# Brief sleep disruption alters synaptic structures among hippocampal and neocortical somatostatin-expressing interneurons

**DOI:** 10.1101/2024.07.22.604591

**Authors:** Frank Raven, Alexis Vega Medina, Kailynn Schmidt, Annie He, Anna A. Vankampen, Vinodh Balendran, Sara J. Aton

## Abstract

**Study objectives:** Brief sleep loss alters cognition and synaptic structures of principal neurons in hippocampus and neocortex. However, while *in vivo* recording and bioinformatic data suggest that inhibitory interneurons are more strongly affected by sleep loss, it is unclear how sleep and sleep deprivation affect interneurons’ synapses. Disruption of the SST+ interneuron population seems to be a critical early sign of neuropathology in Alzheimer’s dementia, schizophrenia, and bipolar disorder - and the risk of developing all three is increased by habitual sleep loss. We aimed to test how the synaptic structures of SST+ interneurons in various brain regions are affected by brief sleep disruption.

**Methods:** We used Brainbow 3.0 to label SST+ interneurons in the dorsal hippocampus, prefrontal cortex, and visual cortex of male *SST-CRE* transgenic mice, then compared synaptic structures in labeled neurons after a 6-h period of *ad lib* sleep, or gentle handling sleep deprivation (SD) starting at lights on.

**Results:** Dendritic spine density among SST+ interneurons in both hippocampus and neocortex was altered in a subregion-specific manner, with increased overall and thin spine density in CA1, dramatic increases in spine volume and surface area in CA3, and small but significant changes (primarily decreases) in spine size in CA1, PFC and V1.

**Conclusions:** Our suggest that the synaptic connectivity of SST+ interneurons is significantly altered in a brain region-specific manner by a few hours of sleep loss. This suggests a cell type-specific mechanism by which sleep loss disrupts cognition and alters excitatory-inhibitory balance in brain networks.

**Significance Statement:** Changes to the function of somatostatin-expressing (SST+) interneurons have been implicated in the etiology of psychiatric and neurological disorders in which both cognition and sleep behavior are affected. Here, we measure the effects of very brief experimental sleep deprivation on synaptic structures of SST+ interneurons in hippocampus and neocortex, in brain structures critical for sleep-dependent memory processing. We find that only six hours of sleep deprivation restructures SST+ interneurons’ dendritic spines, causing widespread and subregion-specific changes to spine density and spine size. These changes have the potential to dramatically alter excitatory-inhibitory balance across these brain networks, leading to cognitive disruptions commonly associated with sleep loss.

## Introduction

Sleep and sleep loss profoundly influence mental health and cognition. As little as a few hours of sleep disruption can alter cognitive performance in both human subjects and animal models ^1–5^. Recent studies have implicated disrupted viability and function of somatostatin-expressing (SST+) interneurons in the early stages of neuropathology driving psychiatric and neurological disorders - including Alzheimer’s disease ^6–10^, bipolar disorder, and schizophrenia ^11–14^. In all these disorders, sleep seems to be disrupted prior to alterations to cognition and behavior ^11–16^.

We and others have recently shown that in mice, short-duration experimental sleep deprivation (SD) in the first few hours following learning experiences can impair both hippocampus-dependent memory processing ^17–24^ and neocortex-dependent memory and synaptic plasticity ^25–31^. How does sleep disruption interfere with these processes, and how might sleep promote them? Recent insights have come from both electrophysiological and biochemical studies of sleep effects on the circuits engaged by information processing, including neocortex and hippocampus. For example, in sensory and motor cortical circuits, post-learning sleep is essential for *de novo* dendritic spine formation ^25^, activation of cellular pathways involved in Hebbian plasticity ^32–34^, and adaptive firing response property changes ^26-30,35,36^.

Numerous studies have suggested that SD may have a particularly disruptive effect on the hippocampal microcircuit ^37,38^. Transcriptomic responses of the hippocampus to SD are numerous and diverse^18,39^. Episodic and spatial memories are easily disrupted by brief SD occurring at either the encoding, consolidation, or recall stages^40,41^. Behavioral memory deficits have been linked to hippocampal subregion-specific disruption of synaptic plasticity ^22–24,42^, principal neuron dendritic spine loss ^22,43^, and neuronal activity disruption^18,19,44–47^.

Much of our understanding of SD effects on brain microcircuitry is based on studies which either used bulk tissue (e.g. transcriptomics or biochemistry), or focused on structure or function of excitatory synapses^22,48^. While much less is known about how inhibitory circuits are affected by SD, recent data suggest that transcriptional and functional responses of inhibitory interneurons to sleep loss can be quite different than those of surrounding principal neurons ^18,19,45^. Moreover, inhibitory synaptic activity varies across sleep-wake cycles, decreasing over wake or brief SD, in both the hippocampus and neocortex ^49–52^. More recent work has suggested that sleep and wake differentially affects the function of synapses corresponding to specific subtypes of interneurons (e.g., SST+ vs. parvalbumin-expressing (PV+) populations) in the hippocampus, with plasticity of the latter, but not the former, being selectively enhanced after a period of wake ^53,54^. Recent work from our lab has demonstrated that SST+ interneurons in the hippocampus are selectively activated by SD, leading to suppression of surrounding principal neuron activity in hippocampal subregions including the dentate gyrus (DG) ^19^. Finally, transcriptomic data from the hippocampus suggests that GABA receptor signaling is dramatically altered by SD in the hours following learning ^18,46^. While it is unclear whether the selective activation of SST+ interneurons by SD has long-term consequences, these data highlight the fact that SD may disrupt cognition via effects on hippocampal interneuron populations ^55^.

SST+ interneurons have recently been shown to be essential for optimal hippocampal memory processing ^19,56^ and neocortical sensory ^57^ and executive functions ^58^. Data from our lab and others suggest that SD-driven changes in the function of neocortical and hippocampal SST+ interneurons may be essential to homeostatic and cognitive responses to sleep loss ^19,54,59^. To clarify how SD affects SST+ interneurons in hippocampus and neocortex, we used recently-developed genetic tools ^60^ to resolve dendritic and synaptic structures in these neurons, and characterized dendritic spine alterations among SST+ populations in different hippocampal and neocortical subregions after a very brief, 6-h interval of SD. Like dendritic spines on principal neurons, available data (e.g. from electron or confocal microscopy with immunohistochemistry) suggest that spines are the major site of synaptic input to SST+ interneurons, and that the majority of the dendritic spines on these interneurons receive glutamatergic synapses ^61–63^. Moreover, these spines appear to be as plastic as those observed principal neurons; additional spines are formed on hippocampal SST+ interneurons’ dendrites in response to learning ^64^. We find that structural alterations are present on SST+ interneurons dendritic spines in every brain region quantified after SD, but the direction, extent, and nature of these changes vary from region to region. The diversity of these changes, and their presence across hippocampus and neocortex, suggests that the inhibitory circuits of the brain are profoundly altered by brief sleep loss. These changes to inhibitory microcircuit elements likely underlie some of the cognitive deficits associated with sleep disruption.

## Materials and Methods

### Mouse husbandry and housing

SST-CRE (*B6N.Cg-Sst*^tm2.1(SST-cre)Zjh^; Jackson) transgenic male mice on a C57Bl6/J genetic background were group housed on a 12 h light/12 h dark cycle (lights on 9 AM – 9 PM) and under constant temperature (22 ± 2°C), with *ad libitum* food and water access, in standard transparent filter-top caging. Prior to behavioral experiments (see below) mice were single housed with beneficial enrichment (i.e., multiple forms of nesting material). SD and sleep monitoring was carried out in each mouse’s home cage. All animal husbandry and experimental procedures were approved by the University of Michigan Institutional Animal Care and Use Committee (IACUC).

### Surgical procedures for AAV-mediated Brainbow labeling

At age 3-15 months (Sleep: 7.1 ± 4.9 months; SD: 6.9 ± 4.0 months) each mouse was transduced with a cocktail of CRE recombinase-dependent AAV vectors to express Brainbow 3.0 (AAV-EF1a-BbTagBY and AAV-EF1a-BbChT; titers ≥ 1×10¹³ vg/mL; Addgene) ^60^ under isoflurane anesthesia. 1 µl of the AAV cocktail was delivered stereotaxically per brain region as follows: to transduce DG, CA3 and CA1 of dorsal hippocampus, injections were placed at: A/P −2mm, M/L ±2 mm, D/V −2.5 mm and −1.5 mm. Coordinates for neocortical areas V1 and mPFC were: A/P −3 mm, M/L ±3 mm, D/V −0.5 mm, and A/P +1.5 mm, M/L ±0.5 mm, D/V −3 mm and −2.5 mm, respectively.

### Handling and SD procedures

Following 9 days of postoperative recovery, mice were single housed and habituated for 5 days to handling procedures (5 min/day). Each mouse was randomly assigned to either Sleep or SD experimental groups. Behavioral experiments started at the beginning of the light phase, two weeks following transduction with Brainbow AAV vectors. SD animals were sleep deprived in their home cage using gentle handling stimulation method for 6 h, during which they were kept awake by gently tapping and/or shaking the cage, and disturbance of nesting material. The necessity of interventions (to disrupt sleep attempts) were documented over the course of the 6-h period, to confirm increases in sleep pressure over the course of deprivation (**Figure 1C**). Mice allowed *ad lib* sleep were monitored visually in 5-min increments to quantify sleep behavior (i.e. assumption of characteristic sleep postures, with closed eyes and immobility) in the home cage (**Figure 1A-B**). This method of sleep quantification has recently been validated as comparable with polysomnographic sleep measurement ^65^. Immediately following the 6-h *ad lib* sleep or SD period, mice were euthanized via an overdose of pentobarbital and were perfused transcardially with ice cold PBS followed by 4% paraformaldehyde for brain tissue harvest.

**Figure 1.**
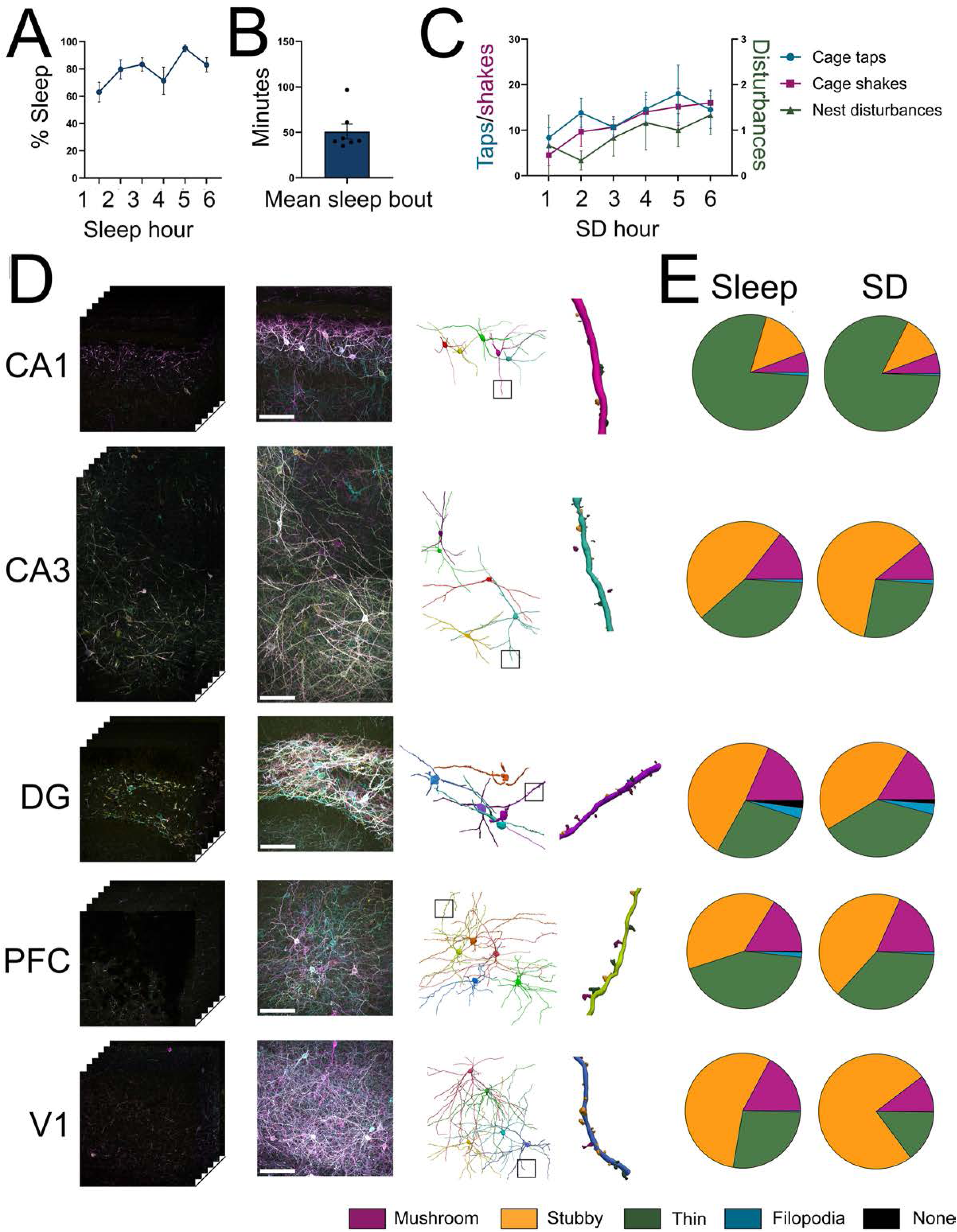
A brief period of SD alters dendritic spine morphology distributions in hippocampal and neocortical SST+ interneurons. Brainbow AAV-transduced SST-CRE mice were sleep-deprived for 6 h via gentle handling (*n* = 6), or were allowed to sleep freely in their home cage (*n* = 7), starting at lights on. *A-B-* Control mice slept for more than 60-70% of the first half of the light phase, with an average sleep bout length of approximately 50 min. *C –* Gentle handling interventions needed to prevent sleep in SD mice (cage taps, cage shakes, and nest disturbances) increased over the course of the 6-h SD experiment. *D-G-*Representative images across the z-axis (**D**), maximum intensity projection images (*E*) showing Brainbow-labeled SST+ interneurons in each of the examined brain regions, corresponding traced neuronal dendritic arbors (*F*), and detail of inset, highlighting dendritic spines (*G*). Scale bars indicate 100 µm. *H -* Distribution of spine subtypes for quantified for neurons in each region in freely-sleeping and SD mice.

### Tissue processing and immunohistochemistry

Perfused brains were post-fixed for 24 hin 4% paraformaldehyde for 24 h at 4°C, washed with PBS, and sectioned coronally (80 µm thickness) using a vibratome. This section thickness was chosen to maximize the extent of dendrite sampling in the Z-plane, without compromising our ability to resolve dendritic spines (i.e., due to light scatter) ^60,66^. Free floating brain sections were blocked overnight at 4°C in PBS containing 5% normalized goat serum and 1% Triton X-100. Each section was then incubated for 72 h at 4°C in PBS with 5% normal donkey serum, 0.5% Triton X-100 and the following primary antibodies chicken anti-EGFP, rat anti-mTFP1, and guinea pig anti-MK2 (all at 1:1000). Following three 1-h rinses at room temperature in PBS with 1% normal donkey serum and 0.5% Triton X-100, sections were incubated for 72 h at 4°C in PBS with 5% normal donkey serum, 0.5% Triton X-100 and the following secondary antibodies: donkey anti-chicken Alexa633, donkey anti-rat Alexa488, and donkey anti-guinea pig Alexa555 (all 1:1000). After secondary antibody incubation, sections received three 1-h rinses and were cover slipped using ProLong Diamond antifade mounting medium (ThermoFisher Scientific).

### Imaging and 3-dimensional reconstruction of dendritic arbors

Images of Brainbow-labeled neurons were acquired using an SP8 confocal microscope (Leica Microsystems) with a 40x oil-immersion objective lens. Images spanned at least 40 µm in depth, using 0.3 µm acquisition steps. Full Z-stack images (0.189 µm/voxel in the X and Y planes, 0.3 µm Z steps) were exported using LAS X software (Leica Microsystems) and traced using Neurolucida360 (MBF Bioscience). Similar results (in terms of numbers of detected spines per dendrite, and their categorization into different spine types) were obtained for images taken at a higher (63x) magnification. For each transduced brain region and animal, Brainbow-labeled neurons with the most complete dendritic arbors represented within the Z-stack (i.e., with no cropping of dendrites within the X and Y planes, and the most complete sampling across the Z plane) were identified. Of those neurons with fully-imaged dendritic arbors, 4-5 were selected at random within each region for 3D reconstruction. Somas were detected automatically (with detection sensitivity varying between 50-80% depending on the signal contrast and intensity), and dendritic arbors were semi-automatically traced. Neurons were selected for analysis with the aim of limiting truncation of dendritic arbors; where dendrites were truncated at a length ≤ 10 µm, these dendrites were excluded from subsequent analysis. Dendritic spines were automatically detected along the full length of traced dendrites, using the following settings: outer range between 1.5 and 3 µm, a minimum height of 0.4 µm, and detector sensitivity set between 75 and 120 µm. Using Neurolucida 360’s preset definitions, spine morphology was classified automatically into one of 4 classes: mushroom, stubby, thin, or filopodium ^67,68^. The criteria for assigning spines to a specific class are based on prior work quantifying morphology of principal neurons ^69^, where spine head diameter/neck diameter ratios, backbone length/head diameter ratios, overall backbone length, and overall head diameter are used for classification. In rare instances, spines could not be classified into any of these 4 classes - these are referred to as “none” in **Figure 1**. These non-classifiable spines were most abundant in the DG hilus, where they may reflect “complex spines” or “thorny excrescences” - morphologically unusual synapses previously characterized within that structure, postsynaptic to mossy fiber terminals ^70,71^. Each spine’s location along the dendrite was used to calculate overall spine density, and for Sholl analysis of spine distributions along the dendrite. Within each class of spine, surface area and volume parameters were used for subsequent analysis. Spine surface area was defined as the apparent exterior surface area of the spine (in µm^2^). Spine volume was calculated as the number of voxels that make up the spine object multiplied by the volume of a single voxel (in µm^3^).

### Statistical analyses of morphological data

Experimenters were blinded to animal treatment conditions throughout image analyses. Data analyses and generation of summary figures were carried out using GraphPad Prism10 (v10.2.3). To examine the distribution of the different spine types *within* a region, and how the distribution of spine types differed *between* regions, a nested one-way ANOVA was used, followed by Tukey’s multiple comparisons tests. For comparisons of spine densities of SST+ interneurons in each brain region between SD and Sleep mice, a nested t-test was used. For Sholl analyses—testing how spine densities vary by distance from the cell soma—we used a repeated-measures two-way ANOVA with a Geisser-Greenhouse correction, using sleep condition and distance from soma as between factors, and subject as within factor. These tests were followed up by multiple comparisons using Fisher’s LSD test, to examine if SD affects spine density at a specific distance from soma, for example. Because dendrite length was variable between neurons, Sholl analysis for each subregion was based on shortest common dendrite length for which measures were available for all reconstructed neurons. Data are presented as mean ± S.E.M. Unpaired Kolmogorov-Smirnov (KS) tests were used to compare the distributions of surface area, volume, spine head diameter, and backbone length of spines between SD and Sleep conditions.

## Results

### Experimental sleep disruption during the rest phase alters the distribution of dendritic spine morphology in SST+ interneurons

To characterize changes to the SST+ interneuron network following sleep deprivation (SD), SST-CRE mice were transduced with AAV vectors to express Brainbow 3.0 ^60^ in a CRE recombinase-dependent manner in prefrontal cortex (PFC), primary visual cortex (V1), and dorsal hippocampal DG, CA3, and CA1. Following postoperative recovery, each mouse was habituated to daily handling for 5 days, and were then randomized into freely-sleeping control (Sleep) and SD conditions. Beginning at lights on, mice were either allowed *ad lib* sleep, or were sleep deprived using gentle handling in their home cage over the next 6 h, after which all mice were perfused for SST+ interneuron imaging and morphological analysis. Sleep behavior was scored behaviorally in control mice, which spent 60-70% of the first 6 h of the rest phase sleeping (**Figure 1A-B**). SD mice showed the expected increase in sleep pressure across the 6-h period of sleep disruption, as evidenced by the increased frequency of interventions (cage taps, cage shakes, next disturbances) needed to maintain wakefulness as SD continued (**Figure 1C**).

Dendritic arbors of Brainbow-labeled SST+ interneurons (*n* = 4-5 neurons per region, per animal) were traced for semi-automated spine detection along their entire length (**Figure 1D**). Within the hippocampus, traced Brainbow-labeled neuronal cell bodies were located in the hilus of DG, in the stratum oriens and radiatum of CA3, and in the stratum oriens of CA1. In neocortical structure (PFC and V1), labeled neurons with cell bodies in layers 2/3 and 5 (including putative Martinotti cells)^72,73^ were traced. Reconstructed dendritic arbors of hippocampal SST+ interneurons were far sparser (particularly in CA1 and CA3) than those traced in neocortical subregions, consistent with previous reports of SST+ interneuron morphology in these regions ^74–76^. Detected spine morphologies were categorized as thin, stubby, mushroom, or filopodial – using classification methods derived from analysis of dendritic spines on excitatory neurons ^69^. Available data from principal (i.e., glutamateric) neurons suggests that stubby and mushroom spine types represent more mature, stable synapses, while thin spines and filopodia are thought to be plastic and/or transient ^67,68^. Across all imaged brain areas, nearly all detected spines on SST+ interneurons’ dendrites could be classified into these same morphological categories – suggesting similar overall dendritic spine structure between SST+ interneurons and principal neurons. Furthermore, across all brain regions measured, the relative distribution of spine types were very comparable to the distribution reported for principal neurons ^9^ – although with generally fewer mushroom spines and a greater proportion of stubby spines ^77,78^. A small percentage (1-2%) of identified spines in DG (and almost none of those identified in other brain regions) had morphological features that did not meet definitions for the four categories; these atypical spines, which are categorized as “none” in **Figure 1E**, are consistent with previously-described “complex spines” or “thorny excrescences” in the DG hilus (where SST+ interneurons’ cell bodies and dendrites reside) ^70,71^. In freely sleeping control mice, thin spines constituted the majority of spines in CA1 and PFC SST+ interneurons, while stubby spines were the predominant spine type in DG, CA3, and V1. SD led to changes in the distribution of the four spine morphologies in hippocampal SST+ interneurons. In both CA1 and DG, SST+ interneurons had a decreased proportion of stubby and mushroom (presumptive mature) spines and a corresponding increase in thin (presumptive plastic) spines after SD. CA3 interneurons showed an increased proportion of stubby spines, with reductions in thin and mushroom spines. Similarly, after SD, the proportion of SST+ interneuron spines with stubby morphology increased in both neocortical areas (PFC and V1), with a corresponding decrease in the proportion of thin spines (and a relative decrease in mushrooms spines in V1). While filopodia were generally rarer in PFC and V1 in freely sleeping mice, the relative proportion of filopodia decreased in both neocortical structures after SD. Thus, if we extrapolate putative spine functions based on morphological types characterized in principal neurons ^67,68^, these data suggest different putative effects of SD on SST+ interneurons’ synapses, depending on brain region. Namely, SD reduces mature and stable spine types among interneurons in CA1 and DG, while simultaneously increasing immature and plastic spine types. In contrast, mature and stable spine types are more numerous in CA3 and neocortical structures after SD, at the expense of immature and transient spine types.

### SD differentially affects overall SST+ interneuron spine density in hippocampal and neocortical subregions

Overall spine density and total dendrite length, but not total spine numbers, for SST+ interneurons varied by brain region (spine density: F_(4, 23)_ = 2.9, *p* = 0.0446; total dendrite length: F_(4, 23)_ = 57.41, *p* < 0.0001; spine numbers: F_(4, 23)_ = 1.4, *p* = 0.2650; **Figure 2A-B**), with dendrites of labeled interneurons in neocortical areas generally longer than those in hippocampus, and the shortest dendrites present in CA1 and DG. Spine numbers on SST+ interneurons appeared to be affected by SD with increases observed after SD on interneurons in CA1 (t = 2.388, df = 9, *p* = 0.0407. This led to significant differences in spine density, with increases in CA1 SST+ interneuron overall spine density (nested t-test; t = 2.473, df = 9, *p* = 0.0354) in SD mice. When densities of distinct spine types were compared separately (**Figure 2D**), SD-driven, type-specific changes were evident in CA1 (where thin spine density increased significantly [ nested t-test; t = 2.992, df = 9, *p* = 0.0152]). Together these data suggest that SST+ interneurons in hippocampus vs. neocortex have divergent responses to SD, with an apparent increase in synaptic input to CA1 interneurons.

**Figure 2.**
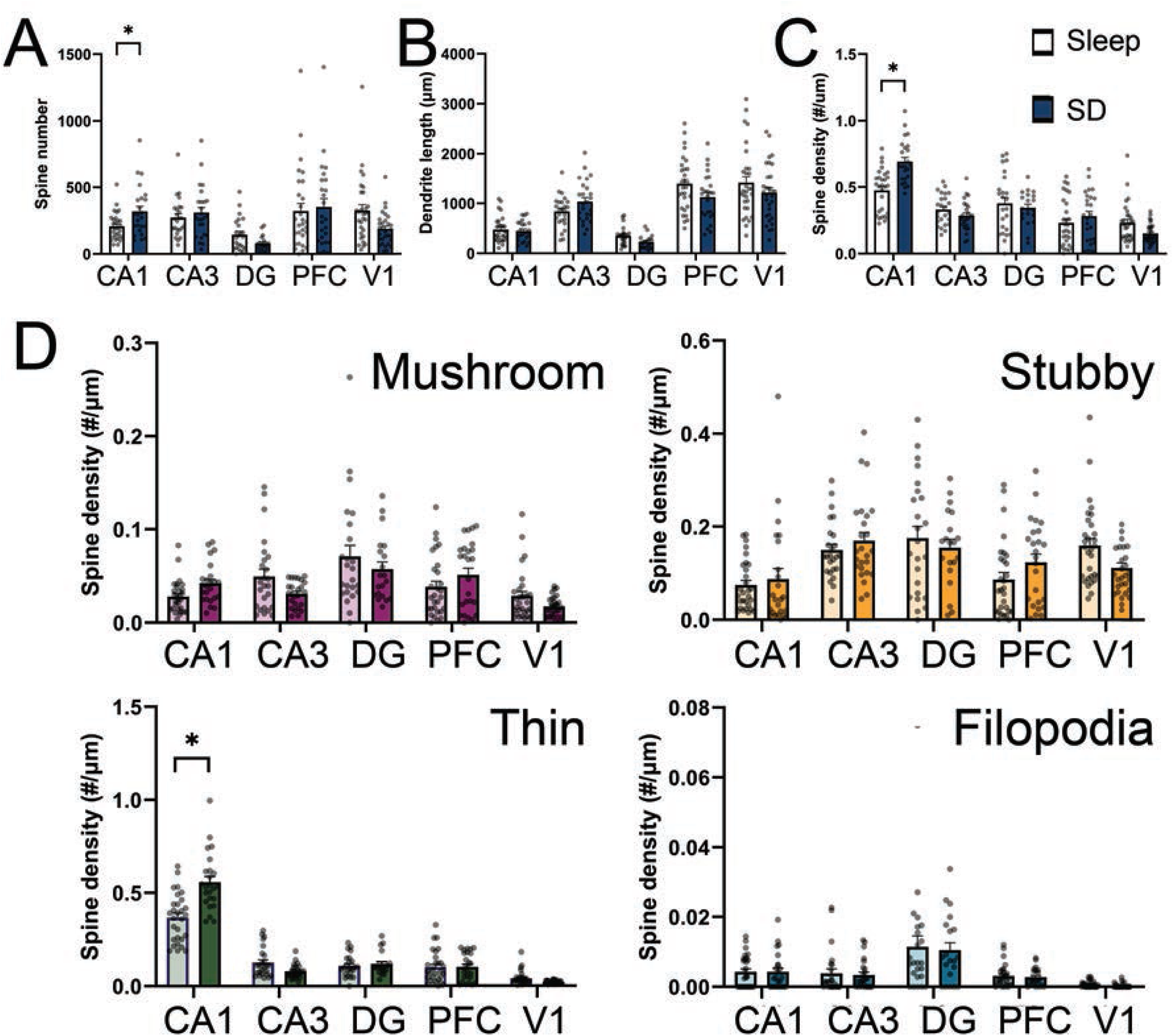
SD differentially affects SST+ interneurons’ spine density in hippocampal and neocortical subregions. ***A-C-*** SD significantly increased total spine density in hippocampal CA1, but did not significantly affect spine density in SST+ interneurons sampled from other brain regions. ***D-G -*** Spine density in each region by type. Thin spine density (***F***) was increased in CA1 after SD. Data points are shown for *n* = 24-30 total neurons/region/condition, with 4-5 neurons/region/animal. * indicates *p* < 0.05, nested t-test.

Prior work has suggested that the SST+ interneuron population is selectively diminished in hippocampus and neocortex with aging, and that loss of these interneurons is a hallmark of age-related cognitive decline ^7,79^. Because mice used in this study covered a range of ages (which were similar in the Sleep and SD conditions), we were interested in whether SD effects on spine density were age-specific. Comparing data from young adult mice (3-5 months of age) with those from older adult mice (10-15 months of age), the same apparent increase in overall spine density was observed in CA1 with SD. There were no apparent age-related changes in overall SST+ interneuron spine density in any of the brain regions sampled.

Because SD is known to alter spine density of CA1 and DG principal neurons at specific distances from their somas ^22,43^, we next tested whether the effects of SD on SST+ interneurons’ spine density were spatially constrained using Sholl analysis. In hippocampal region CA1, which had the most dramatic changes in SST+ interneurons’ total spine density with SD, we found main effects of sleep condition (F_(1, 9)_ = 7.15, *p* < 0.0255) and distance from soma (F_(3.62, 32.6)_ = 7.42, *p* < 0.001), but no significant interaction effect (F_(17, 153)_ = 1.23, *p* < 0.2454). Similar results were found for analysis of CA1 thin spine density (main effect of sleep condition, F_(1, 9)_ = 8.77, *p* = 0.0159; main effect of distance from soma, F_(3.67, 33)_ = 4.52, *p* = 0.0061; sleep × distance interaction, F_(17, 153)_ = 1.20, *p* = 0.2705). While densities of some spine types varied significantly with distance in other brain regions, such as thin and stubby spines in CA1 and CA3, respectively, which gradually increased with distance from soma (CA3 stubby spines main effect of distance from soma: F_(3.65, 29.2)_ = 3.80, *p* < 0.0153), there were no interaction effects where SD affected density as a function of distance. From these data we conclude that SD drives spatially uniform changes in spine density, affecting spines across the length of SST+ interneurons’ dendrites.

### SST+ interneurons’ dendritic spine size and shape change after SD in a subregion-specific manner

Because SD can alter dendritic spine size in principal neurons in both neocortex and hippocampus ^48,80^, we also compared SST+ interneurons’ spine size within each brain region between Sleep and SD mice. We first simply compared ratios of spine volume to surface area, across all spine types, within each brain region. We found that volume-to-surface area ratios were decreased for CA1 SST+ interneurons’ dendritic spines after SD, and increased in CA3 and PFC (*p* < 0.0001, Mann-Whitney test for each region). No significant changes in this ratio for interneurons in DG or V1. Such changes have been linked to plasticity-driven changes in spine biophysical properties, organelle content, and postsynaptic density surface at glutamate receptor-containing spines in principal neurons^81–83^. This suggests that glutamatergic postsynaptic responses in SST+ interneurons different brain regions may be bidirectionally modified by SD.

To resolve these changes in greater detail, cumulative distributions of spine surface area and volume were generated for each morphologically-defined spine subtype and brain region. The most impressive changes in spine size after sleep loss were observed in CA3 interneurons, where the size of all spine types (**Figures 3B** and **4B**) was increased dramatically after SD. Both surface and volume for CA3 interneurons’ stubby (surface area KS test; D = 0.1402; *p* < 0.0001; volume KS test; D = 0.1563; *p* < 0.0001), mushroom (surface area KS test; D = 0.1119; *p* < 0.0001; volume KS test; D = 0.1303; *p* < 0.0001), and thin spine types increased dramatically after SD (surface area KS test; D = 0.1016; *p* < 0.0001; volume KS test; D = 0.1128; *p* < 0.0001). CA3 interneurons’ filopodia, while far fewer in number, also showed significant increases in volume (Kolmogorov-Smirnov test; *p* = 0.0268). Because classification of dendritic spines depends on ratios of spine head diameter, neck diameter, and backbone length ^69^, we also directly compared distributions of head diameter and backbone length (across all detected spines) after Sleep vs. SD. As shown in **Figure 5**, spine head diameters among CA3 interneurons were also dramatically larger after SD (KS test; D = 0.1671; *p* < 0.0001). The ratio of CA3 interneurons’ spine head diameter to backbone length ratios increased overall after SD (*p* < 0.0001, Mann-Whitney test). Together, these data suggest that SD leads to growth and maturation (and potentially, functional strengthening^81–83^) of synapses on SST+ interneuron dendrites in CA3.

**Figure 3.**
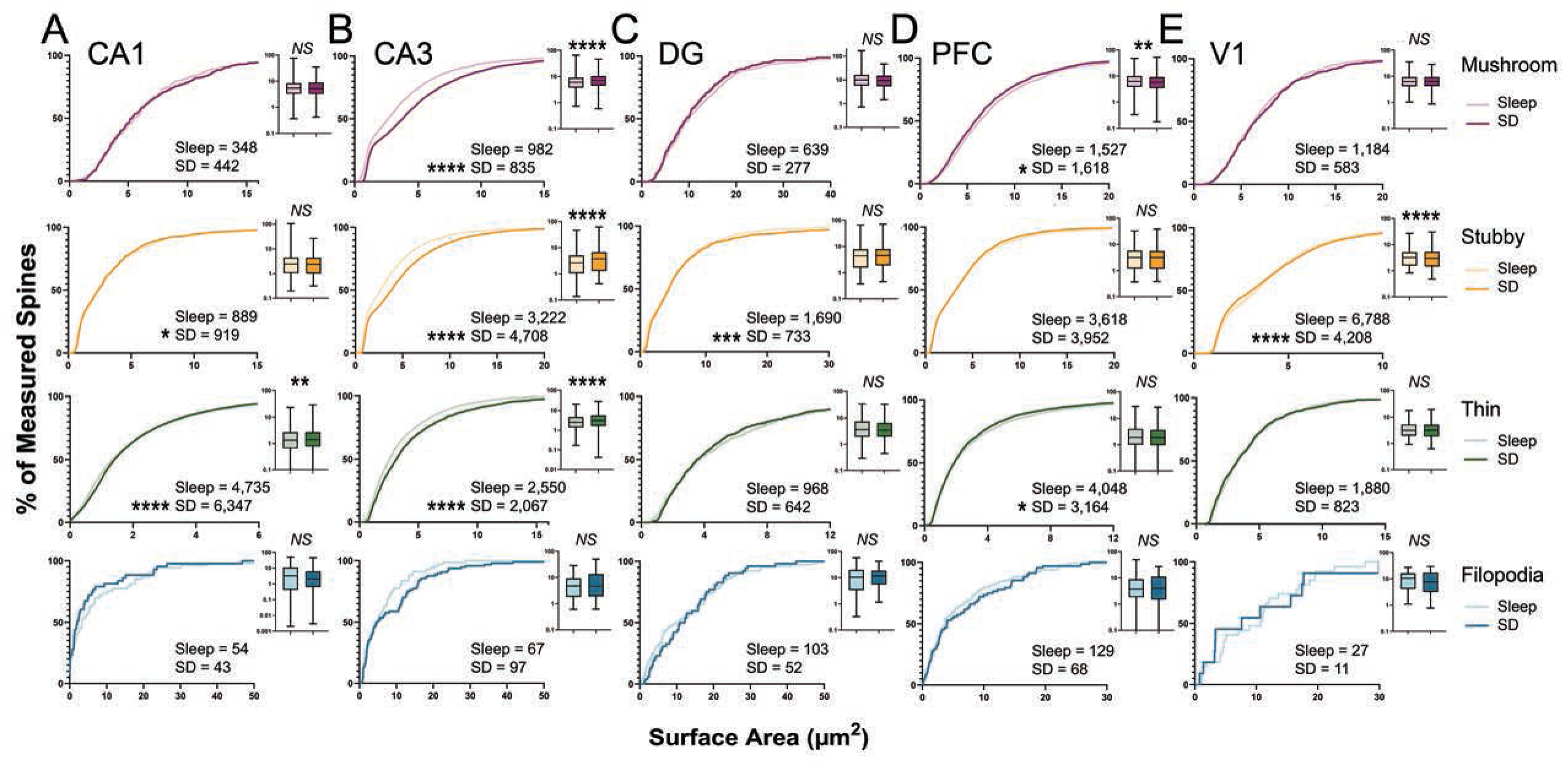
SD alters SST+ interneuron spines’ surface area in a subregion-specific manner. ***A-E -*** SD increased all spine types’ surface area in CA3 except for filopodia, slightly increased stubby and thin spines’ surface area in CA1 and DG, and simultaneously decreased mushroom/thin and stubby spines’ surface area in PFC and V1, respectively. Insets show distributions (25^th^/50^th^/75^th^ percentile, whiskers indicate min and max) of surface area on a log scale (in μm^2^). For insets, ** and **** indicate *p* < 0.01 and *p* < 0.0001, Mann-Whitney test. For cumulative distributions, *, ***, and **** indicate *p* < 0.05, *p* < 0.001, and *p* < 0.0001, K-S test.

**Figure 4.**
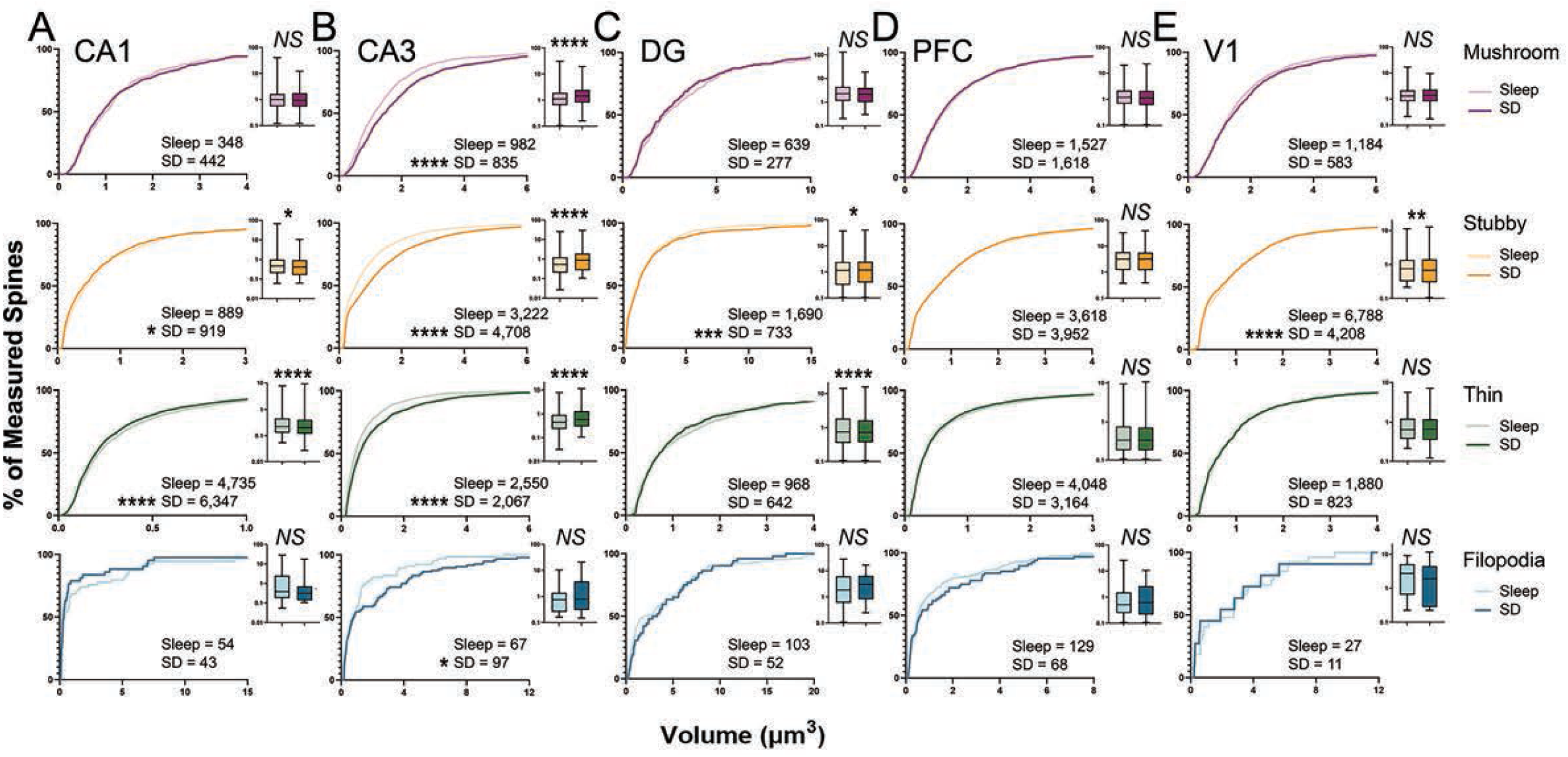
SD alters SST+ interneuron spines’ volume in a subregion-specific manner. ***A-E -*** SD increased all spines types’ volume in CA3, slightly decreased CA1 thin and stubby spines’ volume, slightly increased DG stubby spines’ volume, and decreased stubby spines’ volume in V1. Insets show distributions (25^th^/50^th^/75^th^ percentile, whiskers indicate min and max) of volume on a log scale (in μm^3^). For insets, *, **, and **** indicate *p* < 0.05, *p* < 0.01, and *p* < 0.0001, Mann-Whitney test. For cumulative distributions, *, ***, and **** indicate *p* < 0.05, *p* < 0.001, and *p* < 0.0001, K-S test.

**Figure 5.**
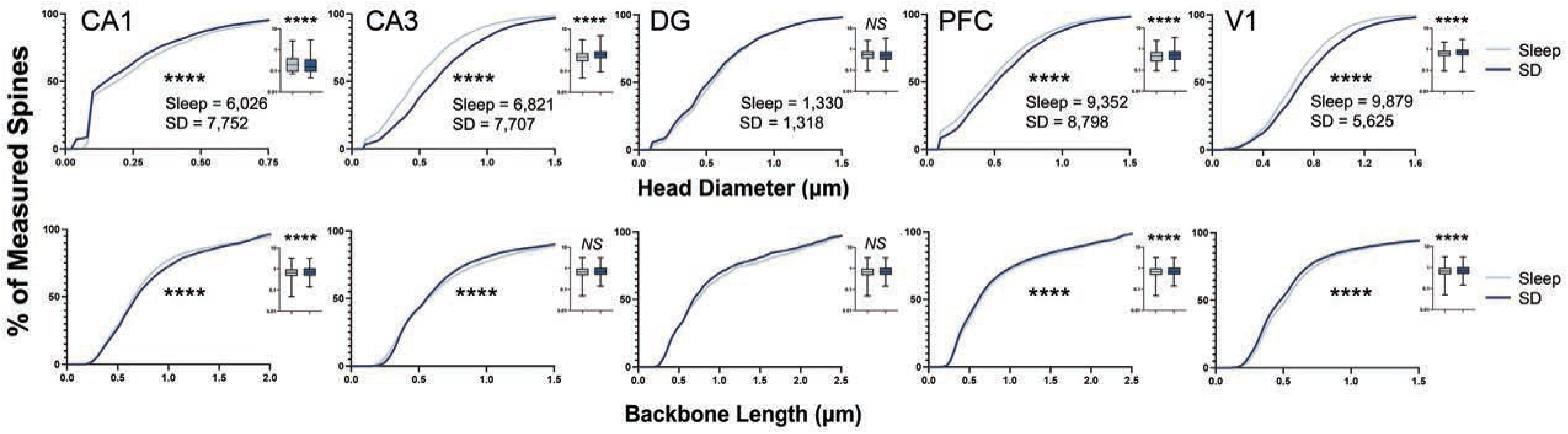
SD differentially affects spine head diameter and spine length in specific brain regions. SD increased spine head diameter in all regions except DG. Spine head backbone length was decreased in the CA1 area, but increased areas CA3, PFC, and V1. Insets show data distributions as described for Figures 3 and **4**. For insets and cumulative distributions, **** indicates *p* < 0.05, *p* < 0.01, and *p* < 0.0001, Mann-Whitney test or K-S test.

In contrast with the dramatic increases in size for all spine types in CA3 interneurons, far more modest changes in spine size were observed after SD for interneurons in CA1 and DG. In CA1, SD had the most pronounced effect on the density of thin spines – the most abundant spine type on CA1 SST+ interneurons’ dendrites (**Figure 1E**). In contrast with the spine size increases observed in CA3, CA1 thin spines’ volume decreased after SD, while their surface area slightly, but significantly, increased (surface area KS test; D = 0.06139; *p* < 0.0001; volume KS test; D = 0.04352; *p* < 0.0001; **Figures 3A** and **4A**). These changes would be consistent with an increase in longer, but thinner, spines on CA1 interneurons’ dendrites. Consistent with this interpretation, spine head diameters (thought to increase with maturation into mushroom-type spines) decreased in CA1 after SD (KS test; D = 0.07856; *p* < 0.0001), while spine length (which is greater in thin spines and filopodia) increased (KS test; D = 0.05087; *p* < 0.0001) (**Figure 5**). Stubby spines - the second most abundant type on CA1 SST+ interneurons - showed small, but significant, decreases in surface area and volume after SD (surface area KS test; D = 0.07236; *p* = 0.0170; volume KS test; D = 0.07301; *p* < 0.0162; **Figures 3A** and **4A**). Overall ratios of spine head diameter to neck length (measured across all detected spines) decreased significantly in CA1 after SD (*p* < 0.0001, Mann-Whitney test). Because both overall and (more specifically) thin spine density is increased in CA1 after SD, these size distribution changes are likely due to new, less-mature spines being added to SST+ interneurons’ dendrites during SD.

In DG, the effects of SD on dendritic spine morphology were minimal – with small, but significant, increases in the size of stubby spines (stubby surface area KS test; D = 0.09781; *p =* 0.0001; stubby volume KS test; D = 0.09399; *p* = 0.0002; **Figures 3C** and **4C**), and no significant changes in other spine types. Overall distributions of spine head diameter (KS test; D = 0.05006; *p* = 0.0724) and backbone length (KS test; D = 0.04447; *p* = 0.1459) were unchanged after SD in DG (**Figure 5**). Taken together, these data suggest that dramatic and wide-ranging changes to interneurons’ synaptic structures take place across the hippocampal network after only a few hours of SD.

Neocortical SST+ interneurons also showed relatively modest changes to spine size after SD. In PFC, SD caused small decreases to mushroom and thin spine surface area (mushroom surface area KS test; D = 0.05752; thin surface area KS test; D = 0.03284; *p* = 0.0434; **Figure 3D**), with no corresponding significant changes to spine volume (**Figure 4D**). Statistically significant morphological changes in V1 were constrained to stubby spines, which showed decreases in surface area and volume after SD (surface area KS test; D = 0.05054; *p* < 0.0001; volume KS test; D = 0.05417; *p* < 0.0001; **Figures 3E** and **4E**). In both neocortical structures, these changes were associated with overall increases in spine head diameter (PFC - KS test; D = 0.06737; *p* < 0.0001, and V1 - KS test; D = 0.09627; *p* < 0.0001) and decreases in spine length (PFC - KS test; D = 0.03527; *p* < 0.0001, and V1 - KS test; D = 0.05266; *p* < 0.0001) after SD (**Figure 5**). In PFC, these changes led to significantly higher spine diameter-to-length ratios (*p* < 0.0001, Mann-Whitney test), while in V1, this ratio was not significantly changed. These changes may be related to the overall reduction in the proportion of thin spines (and increase in stubby spines) we observe in PFC and V1 after SD (**Figure 1**). Taken together, these data suggest that the most pronounced effects of SD among neocortical SST+ interneurons are related to dendritic spine maturation.

## Discussion

Our data present a dynamic picture of how relatively brief (i.e., 6-hour) SD reconfigures inhibitory networks across the dorsal hippocampus and neocortex (**Figure 6**). We find that in hippocampus, where SST+ interneurons are selectively activated by SD ^19^, changes to SST+ interneurons’ dendritic morphology vary widely by subregion. For example, in DG, almost no changes in dendritic spine morphology or density are present after SD. At the same time, SD causes a large increase to the size (but not density) of *all* spines on CA3 SST+ interneurons’ dendrites. Meanwhile, CA1 interneurons’ spine density is greatly increased by SD, with a simultaneous small decrease in stubby and thin spine volumes. Neocortical SST+ interneurons’ structures were also affected by SD, albeit modestly - with some spine types slightly decreased in size in both PFC and V1.

**Figure 6.**
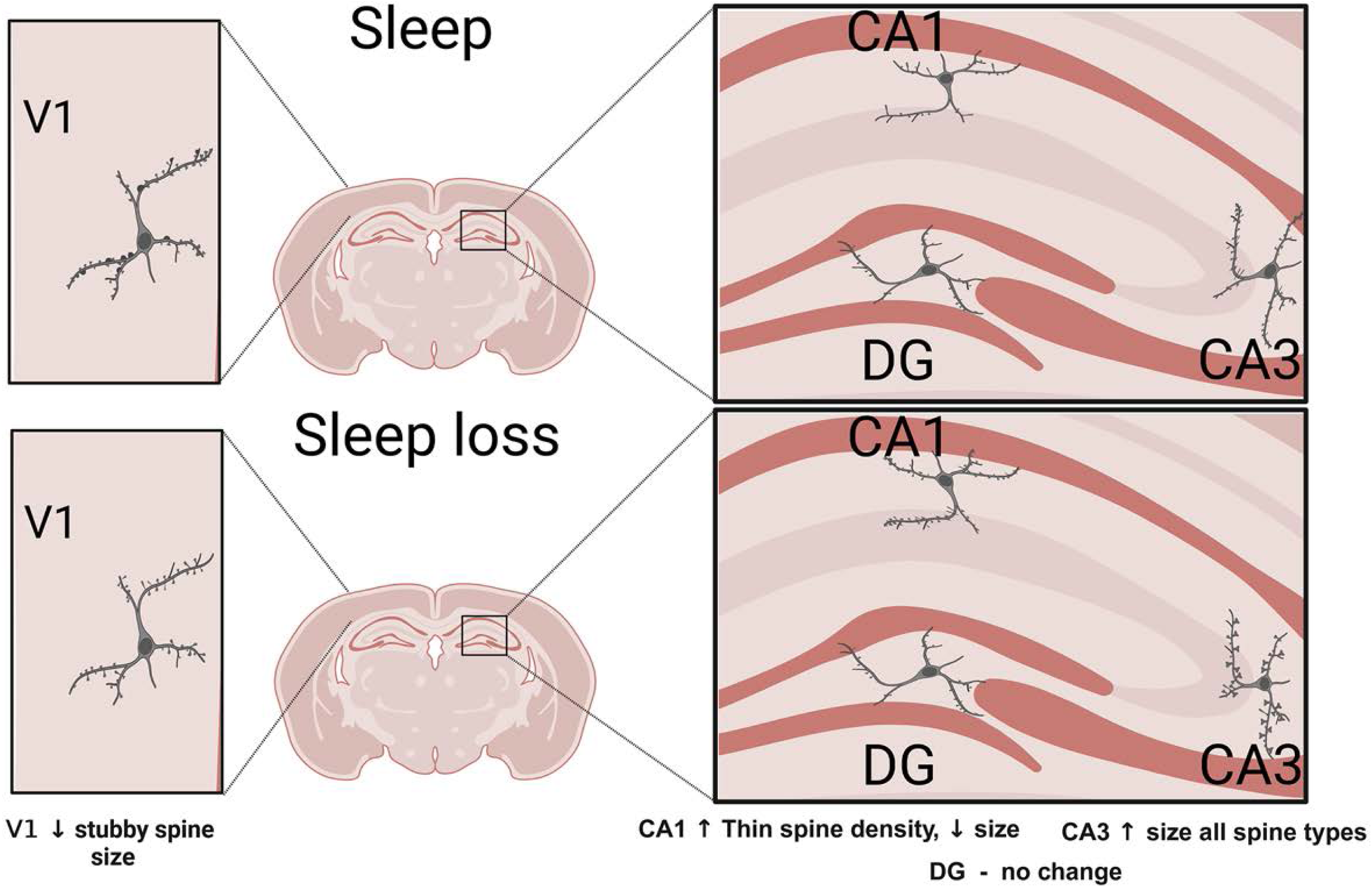
Summary of changes to SST+ interneuron structures in neocortex and hippocampus.

Critically, recent data suggest that spines on SST+ interneurons have similar properties to those on principal neurons. For example, spines associated with these interneurons appear to receive input from glutamatergic and GABAergic presynaptic partners in proportions similar to that observed in pyramidal neurons, with the majority being glutamatergic ^61–63^. Moreover, SST+ interneurons’ spines exhibit similar learning-driven plasticity, e.g. *de novo* spine formation ^64^. Finally, the distributions of spine types we observe on SST+ interneurons’ dendrites (**Figure 1**), and changes in density we observe along the dendrites (**Figure 2**), are similar to those described for pyramidal neurons in hippocampus and neocortex ^9,77,78^. Because available data suggest that spine functions are conserved between SST+ interneurons and better-characterized principal neurons, we can make inferences about how SD affects the function of these interneurons, and the likely consequences for the networks they reside in. For example, increases in spine density in CA1 SST+ interneurons is likely linked to increased excitatory input to, and activity in, this population during SD ^18,19^. In turn, an increase in SST+ interneuron-mediated inhibition could drive overall suppression of activity in the hippocampal network after SD ^17,19,84^, and corresponding dendritic spine elimination in surrounding pyramidal neurons ^22^

Cortical SST+ interneurons (and in particular, Martinotti cells) have been implicated in regulating non-rapid eye movement (NREM) slow wave activity, a marker of the homeostatic sleep response ^59^, and a subset of SST+ interneurons in PFC (a population projecting to hypothalamic sleep regulatory centers) are activated by SD to increase sleep preparatory behaviors and NREM sleep time ^54^. However, the effects of SD on the cortical SST+ population appear to be mixed, with reports of activity increases or decreases reported for different neocortical structures ^54,59^. Here we find that at the level of spine size and shape, SD has similar effects on PFC and V1 SST+ interneurons. In both structures, SD leads to small but significant decreases in spine surface area, volume, and backbone length, and larger increases in spine head diameter (**Figures 3-5**). Within V1, there was also a trend for reduced overall spine density after SD. While available data suggests that postsynaptic responses of V1 SST+ interneurons may be robust to visual deprivation^85^, one possibility is that the increase in visual experience our mice receive during SD could be driving the structural changes we observe. Visually-driven responses in SST+ interneurons tend to be relatively weak and delayed with respect to other V1 neuron populations ^86^, which may make this population susceptible to Hebbian synaptic depression (and associated dendritic spine changes ^87^) in the context of enriched or extensive visual experience. It is worth noting that the morphological spine type most significantly affected (in terms of size and relative proportion) in V1 is stubby spines – a spine type which has been proposed to reflect the retraction of mature (i.e. mushroom) spines as they are eliminated, in the context of synaptic depression ^67,88^. Consistent with this, we find a far greater proportion of stubby spines (at the expense of mushroom and thin spines) after SD in V1 (**Figure 1**). Taken together, our data are consistent with synapse elimination in V1, where overall spine density, stubby spine density, and stubby spine size all show a decreasing trend after a brief period of SD. The overall changes we observe in neocortical SST+ interneurons are consistent with the recent finding (in V1 and PFC) that *ad lib* sleep simultaneously increases synaptic inhibition and decreases synaptic excitation in neocortical networks ^49^. It is also possible that these subtle, SD-driven changes to the structure (and ultimately, function) of these interneurons is directly responsible for changes in slow wave activity observed in the neocortex during recovery sleep ^52,59^.

We identified more complex changes to the SST+ interneuron network within the hippocampus after SD. In DG, where we had previously characterized SD-driven selective activation of SST+ interneurons ^19^, SD had the most minor structural effects – i.e., no changes in dendritic spine density, and a mix of changes to spine size (decreases in thin and mushroom spines, increases in stubby spines). In CA3, spines’ sizes increased dramatically, across all types. In CA1, overall spine density (and density of the most abundant spine type in CA1 – thin spines) increased after SD, but spine sizes tended to slightly decrease. One possibility is that with respect to presumptive excitatory input at these dendritic spines, opposing changes in spine size and density (and opposing changes between regions CA1 and CA3) might represent cellular or circuit-level homeostasis, preserving overall input to individual SST+ interneurons, constraining their firing rate, and maintaining the relative excitatory-inhibitory balance of the network.

The precise time course of these changes, and the specific roles of NREM and REM sleep, could only be answered in future studies using *in vivo* imaging of dendritic spines. Such studies would also provide a more precise measurement of dendritic spines – which almost certainly undergo subtle changes in size during tissue fixation. Critically, both NREM and REM sleep have been shown to affect rates of spine formation and elimination in neocortical pyramidal neurons ^25,89^, and to impact the overall activity of both SST+ and parvalbumin-expressing interneurons ^51,52^ *in vivo*.

An unanswered question is how these changes interact with other functional and structural changes occurring in the brain across SD, and how these changes, together, affect cognitive processes. Recent data from our laboratory have demonstrated that SD drives selective activation in dorsal hippocampal SST+ interneurons, which suppresses activity in surrounding principal neurons ^19^. Converging evidence from cell type-specific translating ribosome affinity purification (TRAP)-seq and spatial transcriptomic studies suggests that modulation of excitatory-inhibitory balance in structures such as DG accompany post-learning SD, which may provide a mechanism for disruption of memory consolidation ^18,19,46^. At the same time, SD can reduce pyramidal neurons’ dendritic spine density in DG ^43^ and CA1 ^22^, and can impair Hebbian plasticity at glutamatergic synapses ^22,24^. Together, these data suggest a mechanism by which SD-driven increased network inhibition, decreased principal cell activity, or both lead to suppression of hippocampal network output. This idea is supported by human neuroimaging studies in which hippocampal memory encoding and associated hippocampal network activation are impaired after a night of SD ^84^. Whether these effects are entirely driven by plastic changes in SST+ interneurons is unclear. However, it is worth noting that SST+ interneurons play a critical role in gating excitatory drive to CA1 pyramidal cells from different input pathways ^90^, and the information encoding capabilities of DG granule cells ^56,91^. They also directly encode information about space and context essential for hippocampal memory, via plastic changes to their excitatory synapses ^71,92–94^. Thus, insofar as the structural changes we observe in hippocampal SST+ interneurons are likely to: 1) outlast the period of SD itself, and 2) disrupt or alter subsequent learning- or recall-driven synaptic interactions or activities in the SST+ population, it is plausible that these changes directly interfere with hippocampal memory processing.

One of the more interesting findings from this study are the differences in SD effects on the SST+ populations in hippocampus vs. neocortex, with the hippocampal SST+ interneuron network undergoing dramatic changes, and neocortical interneurons being only modestly affected. Why might the effects of SD on the hippocampal and neocortical networks differ so dramatically? While neocortical and hippocampal SST+ interneurons universally receive excitatory input from, and provide inhibitory input to, surrounding principal neurons, interneurons expressing SST have remarkable diversity ^95^. This diversity likely extends to function and plasticity. For example, we have previously shown that SST+ interneurons in the hippocampus are selectively activated by SD ^19^, and that this contributes to selective, early, downregulation of overall hippocampal network activity when sleep has been disrupted ^19,55,96,97^. However, there is no present data suggesting that neocortical SST+ interneurons behave similarly in the context of SD, despite their reported roles in regulating sleep homeostatic responses and NREM slow wave activity ^54,59^. Differences in cellular activity and structural plasticity between the structures are likely due to the different developmental ^98^ and transcriptomic ^95,99,100^ profiles of hippocampal and neocortical SST+ interneurons. Even within these structures, multiple subclasses of SST+ interneurons with distinct electrophysiological, anatomical, and transcriptomic properties have been identified. These region- and subclass-specific properties lead to differential responsiveness to features of SD like wake-associated neuromodulator release ^19^, which could in turn impact the plastic changes SST+ interneurons undergo in the context of SD. Future studies will be needed to determine how experiences in wake (and across SD) contributes to these interneurons’ function, and their role in regulating principal neurons’ activity ^101^. However, recent studies have shown that SST+ interneurons in DG and CA3 directly modulate hippocampal network oscillations (e.g. sharp wave ripples, gamma oscillations), and gate corticohippocampal input from the entorhinal cortex ^102^. Thus, the dramatic SD-driven changes to synapses observed in CA3 interneurons likely has important consequences for subsequent hippocampal information processing.

Sleep loss is a growing health concern, affecting millions of people worldwide across all ages, and severely impacting cognitive functioning. The data presented here highlight the diverse and, in some cases, dramatic changes to SST+ interneurons’ structure that a few hours of sleep loss can cause. Changes to SST+ interneurons’ function, and related changes in excitatory-inhibitory balance, have been implicated in the progression of several neurodegenerative and neuropsychiatric disorders, such as Alzheimer’s disease, schizophrenia, bipolar disorder, and major depressive disorder ^6–14,103,104^. Understanding the mechanisms through which sleep loss alters the morphology, connectivity, and function of this interneuron population will have wide implications for treating cognitive disruption in these disorders – where alterations in sleep are a common feature.

## Author contributions

F.R. and S.J.A. conceived and designed the study. F.R., A.V.M., K.S., A.H., A.A.V., V.B., and S.J.A. performed the research. F.R., A.V.M., K.S., A.H., A.A.V., and S.J.A. analyzed the data. F.R. and S.J.A. interpreted the results and wrote the manuscript. S.J.A. supervised the study.

## Acknowledgements

Figures summarizing results was created using BioRender. This work was supported by NIH research grants R01NS118440 and R01MH135565 and a Chan Zuckerberg Initiative Collaborative Pairs Grant to S.J.A.

## Data Availability Statement

All data will be made available to investigators after publication, if requested.

